# Successful microbial colonization of space using an anti-aggregation strategy

**DOI:** 10.1101/2021.07.09.451734

**Authors:** Xiaonan Liu, Miaoxiao Wang, Yong Nie, Xiao-Lei Wu

## Abstract

Many organisms live in habitats with limited nutrients or space, competition for these resources is ubiquitous. Although spatial factors related to population’s manner of colonizing space influences its success in spatial competition, what these factors are and to what extent they influence the outcome remains under-explored. Here, we applied a simulated competitive model to explore the spatial factors affecting outcomes of competition for space. By quantifying spatial factors using ‘Space Accessibility’, we show that colonizing space in an anti-aggregation manner contributes to microbial competitive success. We also find that the competitive edge derived from being anti-aggregation in colonizing space, which results in a higher ‘Space Accessibility’, neutralizes the disadvantage arising from either lower growth rate or lower initial abundance. These findings shed light on the role of space colonization manners on maintaining biodiversity within ecosystems and provide novel insights critical for understanding how competition for space drives evolutionary innovation.

## Introduction

Competition is a ubiquitous phenomenon observed for both microorganisms and macro-organisms(Oliveira *et al.* 2015; Si *et al.* 2019). It is considered to represent a key factor driving biodiversity(Maynard *et al.* 2017; Smith *et al.* 2018) and evolution(Baalen & Yamauchi 2019; Bernhardt *et al.* 2020). Competition often occurs when individuals compete for identical resources(Lloyd & Allen 2015; RJ & DJ 2015; Burson *et al.* 2018; Paquette *et al.* 2018). It is characterized by consumption of a limiting resource by one population, resulting in a decrease in fitness of its competitors. Nutrient and space are the main two resources necessary but usually limited for organisms, and they are tightly related to each other. A population colonizing more space will commonly obtain more nutrients and energy to support their growth(Benayahu & Loya 1981; Lirman 2001; Paquette *et al.* 2018). One population expands into the available free space, and competes with other populations to occupy areas where nutrients are more abundant. Especially for microorganisms, which often settle on surfaces and form a dense biofilm, where nutrient and space limitation is common, and thus results in strong microbial competition.

To win in a game of microbial competition, numerous competitive strategies have been evolved by microorganisms. For example, microbes gain competitive advantages by privatizing nutrient resources. Siderophores(Ali & Vidhale 2013), compounds that bind and transport iron, have long been recognized as public goods. However, certain strains produce privatized siderophores, which are used to outcompete other strains(Niehus *et al.* 2017; Eickhoff & Bassler 2020). Bacteria such as *Dietzia* sp release membrane vesicles binding extracellular iron, presumably to be recruited by closely related species, implying the roles of membrane vesicles in the microbial competition(Wang *et al.* 2021b, c). In addition, microbes obtain fitness benefits by diversifying metabolic mode. A long-term evolutionary experiment using *E. coli* showed that a citrate-using mutant appeared from the starting population, and found that this mutant was able to uptake citrate at highly increased rates. This feature was accompanied by an increased fitness compared with its ancestor strain(Blount *et al.* 2012). *Saccharomyces cerevisiae* was reported to switch its mode of metabolism between fermentation and respiration depending on the external conditions, thus ensuring its fitness is higher than that of competing yeast cells within the vicinity(MacLean & Gudelj 2006). Furthermore, motility and adhesion also help microorganisms win the competition. Motile bacteria accessed new spaces faster than non-motile bacteria(P & K 1981; Scharf *et al.* 2016; Wang *et al.* 2021a), while matrix-secreting cells are highly adhesive and can colonize the free space more stable than non-secreting cells, thus avoid sloughing by shear forces(Petrova & Sauer 2012; Schluter *et al.* 2015). All the competitive strategies above that these organisms have evolved are biotic factors that affect the outcome of microbial competition.

In addition to the biotic factors, microbial competition is also affected by abiotic factors. Several reports have shown that certain abiotic factors, such as temperature and pH, influence the outcome of microbial competition by changing the intrinsic properties of organisms like growth rates. For example, when temperature is set to above the optimum temperature for fast-growing strains, slow-growing strains are often favored, thus dominating the multispecies community(Lax *et al.* 2020). Another study showed that pH changes create feedback loops that can either facilitate or inhibit the growth of bacterial populations and ultimately affect competition outcomes(Ratzke & Gore 2018).

A recent study showed that emigration rates, i.e. rates at which individuals of a population depart from a particular community, influence the outcome of microbial competition; without, however, changing the fitness of the competing organisms. If faster-growing species is strongly inhibited by slower-growing species, slower grower would dominate under low mortality while faster grower would dominate under high mortality (Abreu *et al.* 2019). However, whether there are other abiotic factors that does not influence the population growth rates but will also affect the outcome of microbial competition, remains to be elucidated. Understanding this question is critical to explain how those slow-growing microbes compete against their fast-growing counter-partners and exist in all environments(Gause 1932; Davis *et al.* 2011; Henson *et al.* 2018).

Ecological processes occur not only in time but also in space. Abiotic spatially related factors may also be potential factors independent of biological intrinsic properties but affecting outcomes of the competition for space. Our previous study indicated that even if the initial abundance and inherent fitness of two populations were identical, outcomes of the spatial competition were not completely random but significantly influenced by the relative positions and time orders for the emergence of different genotypes(Wang *et al.* 2020). These findings suggested that specific events that occurred during the space colonization affected which population colonized more space. However, what these events are and how they affect the outcome of spatial competition, has not been studied enough in our previous research, then more research is needed.

Microbial spatial competition is very similar to the traditional Chinese board game, *Go*, in which two players compete for occupying more spaces on a board. In the *Go* game, the players’ strategies in layout and middle stage are crucial to gaining more territory and winning the game(Baker 2008). Therefore, we hypothesized that factors related the manner of microorganisms exploring and colonizing free space, such as initial spatial positions and the subsequent directions of expansion, play a significant role in their competitive outcome.

In this study, we constructed an individual-based model (named “BacGo”) to simulate two microbial populations competing for limiting space and explore the influence of spatially related factors on competition for space. Our work provides a quantitative view of how the manner in which a microbial strain colonizes new spaces affects the outcome of competing with other strains.

## Materials and methods

### Basic settings of the BacGo model

To simulate the spatial competition between two populations, the BacGo model was built based on 2D lattices, following our previous framework(DK 1998; Kreft *et al.* 1998; Wang *et al.* 2020). In our model, a microhabitat was conceptualized as a 20×20 array. One microbial individual was allowed to occupy a specific spatial grid box. Two populations were assumed to compete for this ‘microhabitat’, where they were allowed to grow and reproduce. For simplicity, nutrients are assumed to be unrestricted in the model and the growth rate of each cell was assumed to be constant (*μ*_*max*_), then

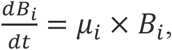

where *B_i_* is the biomass of the *i*th individual; *μ*_*i*_ is the growth rate of the *i*th individual.

In the basic model, the *μ*_*i*_ was set to be same for all cells of both populations, of which the default value was 0.1 fg/fg∙min(DK 1998). The initial biomass of each individual was set as 150 fg(DK 1998). After enough cycles for biomass accumulation, one cell reproduced when its biomass reached the upper threshold of 2*B_0_*. After cell division, the mother cell stayed in the original grid box, while the daughter cell randomly selected one of the 8 directly adjacent grids. If the selected grid has been occupied, the newborn cell will compete for the grid with its aborigine and have a 50% probability to survive. In addition, a randomly death events were considered and set at a very low probability of 1e^−6^. All variables and parameters used in the model were listed in Table S3, the defined indexes were summarized in Table S4, and the abbreviations were summarized in Table S5.

### Simulation protocols and data recording

To simulate the processes of the two populations competing for the ‘checkerboard’, time-lapse numerical simulations lasted for at least 50000 steps, until one population fully occupy the whole space. The model was implemented by C++ language, and the source code are available on https://github.com/Neina-0830/BacGo-model. Simulations were run on an Ali cloud server running Windows Server 2019. Custom functions were included in the Codes to record the position coordinates of every cell, biomass of every cell, as well as the relative abundance of each population at each time point. These simulation data were analyzed and visualized using custom Wolfram Mathematica scripts (https://github.com/Neina-0830/BacGo-model). In particular, in each simulation, the relative abundance of each population at ‘full-occupied’ time point (t_2_) were extracted, and the abundance asymmetry index, AbunR, was calculated as follows:

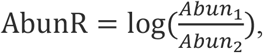

where *Abun*_1_ is the relative abundance of the focus population at t_2_ and *Abun*_2_ is the relative abundance of its competitor at t_2_. AbunR of a population greater than zero means that this population had a higher relative abundance than its competitor at t_2_.

When applicable, 100 replicated simulations were performed for one initial cell distribution, and competition outcomes were summarized to get the winning frequency of the both populations. Then, the winning asymmetry index, WinR, was calculated as follows:

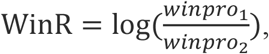

where *winpro*_1_ is the winning frequency of a population in the 100 replicated simulations starting from the same initial distribution, while *winpro*_2_ is the winning frequency of its competitor. When WinR of a population is positive, this population has a higher winning probability than its competitor.

### Parameters describing different colonization manners on spatial competition

To investigate the effect of different colonization manners on spatial competition, we defined serval parameters quantifying spatially factors. To characterize the initial population distributions, a non-dimensional parameter, ScatR, was defined to assess the asymmetry of the scatter level of the initial cell distribution between a population and its competitor, calculated by

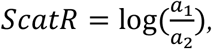

where *a*_1_ and *a*_2_ are the average Euclidean distance between cells of two populations respectively, which characterize the scattered level of initial cell distribution of each population. The ScatR greater than 0 indicates that the population is initially distributed more scatteredly than its competitor, and the absolute value of ScatR represents the degree of the difference in the scatter level of the initial cell distribution between the two populations.

To capture the random events occurring during population expansion in the “occupation stage”, a parameter FreeR was defined to characterize the difference in the degree of ‘expansion freedom’ between one population and its competitor, given by

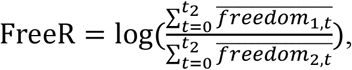

where 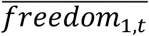 is the average number of empty grids around the daughter cells born in time point *t* of the focus population, while 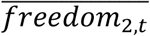 is that number of its competitor. The FreeR index greater than zero indicates that the population possesses greater ‘expansion freedom’ against its competitor in the given simulation, and the higher absolute value suggests a higher difference in expansion freedom between the two populations.

In order to integrate the effect of initial cell distribution and ‘expansion freedom’, a new parameter, named ‘Space Accessibility’, was defined. The asymmetry of ‘Space Accessibility’, SAR, evaluated the competitive edge derived from ‘Space Accessibility’ of the population across whole “occupation stage”, given by

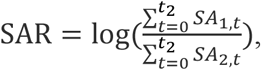

Where *SA*_1,*t*_ and *SA*_2,*t*_ are the ‘Space Accessibility’ for the focal population and its competitor at time point t, respectively. SAR greater than 0 means that the population generally possess higher ‘Space Accessibility’ than its competitor.

### Competition between ‘SmartBac’ and‘NormalBac’

To test the effect of space colonization manners from another perspective, one population was defined to be “smart population” (named as ‘SmartBac’ thereafter), whose daughter cells were always able to colonize the space to ensure that the whole population retained optimal spatial distribution with higher ‘Space Accessibility’. To achieve this goal, ‘SmartBac’ was controlled to colonize space in an anti-aggregation manner. After each daughter cell was born, every possible scenario for its follow-up move were assessed by calculating the *SA*_*k,t*_ value of the formed cell distribution. Then the distribution with maximum *SA*_*k,t*_ value was selected as the next colonizing step of the ‘SmartBac’. To test whether ‘SmartBac’ behaves better in spatial competition, individual-based simulations were performed by considering a competition process between ‘SmartBac’ and ‘NormalBac’ (a normal population), who possessed purely random manner of colonization of space same as the definition in the basic model. This simulation was implemented using a modified C++ code, which is available online on https://github.com/Neina-0830/BacGo-model.

More details about parameters definition and simulations design are given in Supplementary information S1.

### Statistical analysis

The chi-square test was carried out using the *chisq.test* function in stats package of R 4.0.2. Unless indicated otherwise, unpaired, two-tailed, two-sample Student’s t-test was performed for comparative statistics using *t.test* function in stats package of R 4.0.2. Linear correlation analyses between different parameters were implemented using *lm* function in stats package of R 4.0.2. The Cohens’D statistic was calculated using *cohensD* function in lsr package of R 4.0.2. The multiple regression analysis and multicollinearity test were performed using IBM SPSS Statistics 27.0.

## Results

### Simulating competition for space using “BacGo” model

To investigate how spatial positioning of populations affects the outcome of microbial competition, we simulated two populations competing for space with a limiting size by building an individual-based model (named “BacGo”). The model was implemented in discrete grid boxes of a 20×20 array. As shown in Fig. 1a, our simulations were based on three basic assumptions. First, the two competing populations possess the same inherent growth rate and equal initial cell numbers, thus the only differences between them are their manners of colonizing free space. Second, the newly born daughter cell is located around its mother cell but with a random direction of spatial positioning(Kreft *et al.* 1998), resulted in a microcolony with different spatial patterning (Fig. 1a). Lastly, if the selected box has been occupied, the newborn cell will compete for the box against the original occupants of the box and possesses a probability of 50 % to survive(Crowley *et al.* 2004).

**Fig. 1.**
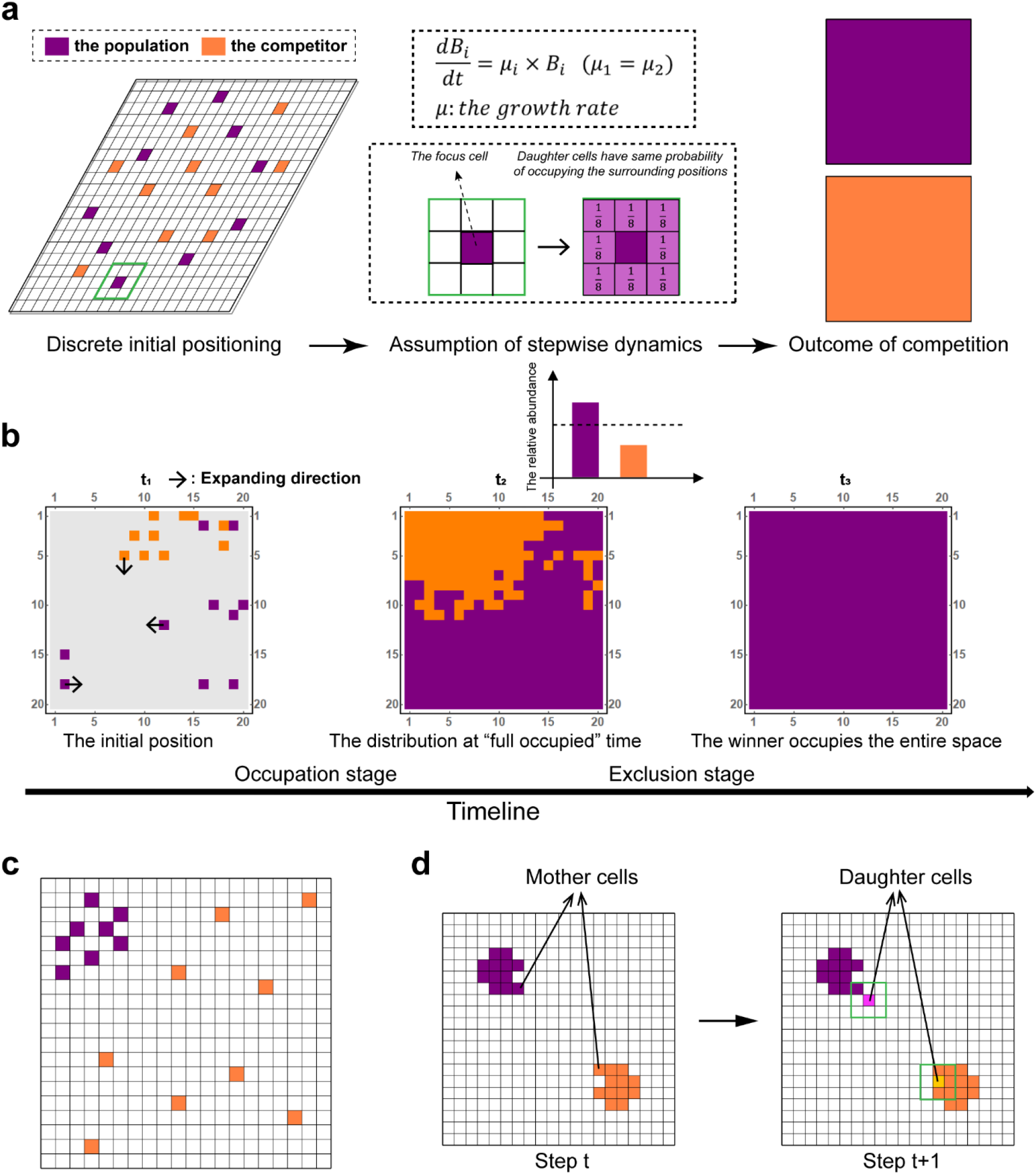
Logic and basic assumptions of the model. **a.** Overall framework of the model. We considered two populations competing for a limited 2D space. The space was initialized by a 1:1 mix of cells from two populations, which were then randomly scattered in the 2D panel. Cells from the two populations exhibited the same growth rate. Cell division occurred when its biomass reached a threshold, during which the daughter cell randomly selected one of the 8 directly adjacent grids. If the selected grid had been colonized, the newborn cell competed for the grid with its original occupant and have a 50% probability to survive. We aimed to see which population ultimately colonized the entire space, and for this purpose investigated the relationship between colonization manner of one population and its competitive success. **b.** The process of spatial competition can be divided into two stages, namely ‘occupation stage’ and ‘exclusion stage’. Small black arrows indicate the direction of population exploitation; t1, t_2_ and t_3_ refer to the initial time point, the “full occupied” time point and the winner colonizes the entire space time point in the process of competitive interaction, respectively. **c.** Representative snapshot shows the difference in the initial distribution between the two populations. The orange population was more scattered than the purple one. **d.** Representative snapshot showing the difference in ‘expansion freedom’ between the two populations. Daughter cells (labelled as light purple) of the purple population were characterized by a higher degree of expansion freedom than the daughter cells (labeled as light orange) of the orange population.

We first explored the outcome of spatial competition, which started by randomly distributing two populations on the grids with the same initial cell numbers of 10 for each. Based on our basic assumptions and the predictions of competitive exclusion theory(Denboer 1986), we hypothesized that only one population could win the competition and finally occupy all grids. As shown in 20000 independent simulations with randomly initial distributions, we discovered that at the end of each simulation, only one population survived. Our Chi-square test results showed no significant difference (P =0.211) between the simulated winning times (10177 of 20000 simulations) and the random winning times (10051 of 20000 simulations) of the two populations. This result conformed with our initial assumption that cells possess a probability of 50 % to survive in the competing with original occupants. When we replicated the simulations initiated with the same cell distributions, we found that the winning probabilities for each population changed in line with the initial distributions (Fig. S1). However, the winning probabilities never reached 100% no matter how the initial distribution changes. Together, these results suggested that yet unknown random factors may affect the final outcome of the competition.

Next, we analyzed the dynamics of microbial colonization during our simulations. As summarized in Fig. 1b, we divided the competition processes into two stages. In the first stage (named “occupation stage”), the cells grew, divided, and occupied the space from the initial positions at time point t1, until the space was fully occupied by both populations at time point tS_2_. In the second stage (named “exclusion stage”), cells from both populations competitively excluded each other, until one population completely filled the entire space (named ‘winner’) at time point t_3_ (Fig. 1b). To statistically characterize the competitive outcome at t_3_, we defined the winning asymmetry index, WinR, of a population as the difference in winning frequency between a population and its competitor in 100 replicated simulations, starting from the same initial distribution. In addition, we defined the abundance asymmetry index, AbunR, of a population as the difference in cell numbers at t_2_ between a population and its competitor. As shown in Fig. S2a, we found a strong positive correlation between AbunR and WinR (R^2^=0.740, P<0.001), indicating that if any one population is more abundant at the “full occupied” time (t_2_), it is more likely to finally win the competition (i.e., occupy the whole space at t3) (Fig. S2b). These results strongly suggest that one population may obtain an asymmetric benefit from the random manners of colonizing space in the “occupation stage”, a benefit that assists this population in colonizing more space at t_2_, thus largely determining the ultimate outcome of the competition.

All of these initial explorations of the model indicated that, in addition to the growth rate(Sebens 1982) and initial cell numbers(Campbell *et al.* 2011), the random manners of colonizing space, such as positioning individuals at initial time and choosing direction of cells during population expansion, provides a considerable competitive edge for a population to colonize a space.

### ‘Space Accessibility’ affects outcomes of spatial competition

#### Larger initial distance of cells is conducive to success in competition

We next investigated what manners of colonizing space determine spatial competition outcomes. Since the competition outcome changes with different initial cell distributions (Fig. S1), we first explored how the differences in the features of initial cell distributions affect the outcome of subsequent spatial competition. Our model assumed that the direct competition between different cells occurred only when cells are located adjacent to each other (assumption 2 and assumption 3). Based on these assumptions, if cells from one population possess greater distance among each other (in other words, distributed more scatteredly), the undesirable intrapopulation competition can be avoided. In addition, space colonization will start from a higher number of seeding positions, thus they may possess higher probability to occupy more free spaces. Therefore, we hypothesized that if one population exhibited a higher degree of scatter across new space to be occupied at time point t_1_, it will potentially occupy more space at time point t_2_, resulting in a higher probability to emerge as the winner.

To compare the levels of scatter (Fig. 1c) of initial cell distribution between two populations, we defined the scatter asymmetry index, ScatR, of a population as the difference in the average distance among different cells at t_1_ between a population and its competitor. To investigate whether the initial scatter level affects the competition outcome, we selected 215 initial cell distributions randomly, which covered a gradient of ScatR values ranging from −1.053 to 1.053 (Blue lines in Fig. S3). We then performed 100 replicated simulations for each initial distribution, to reveal the relationship between the competition outcome and ScatR. Our results showed that AbunR is positively associated with ScatR at significant levels (Fig. 2a; R^2^=0.284, P<0.001), indicating that the population initialized with more scattered cell distribution will occupy more space at t2. Moreover, a positive relationship was also observed between WinR and ScatR (Fig. 2b; R^2^=0.324, P<0.001), further suggesting that the benefit obtained from more scattered initial cell distribution contributes to the ultimate dominance of this population.

**Fig. 2.**
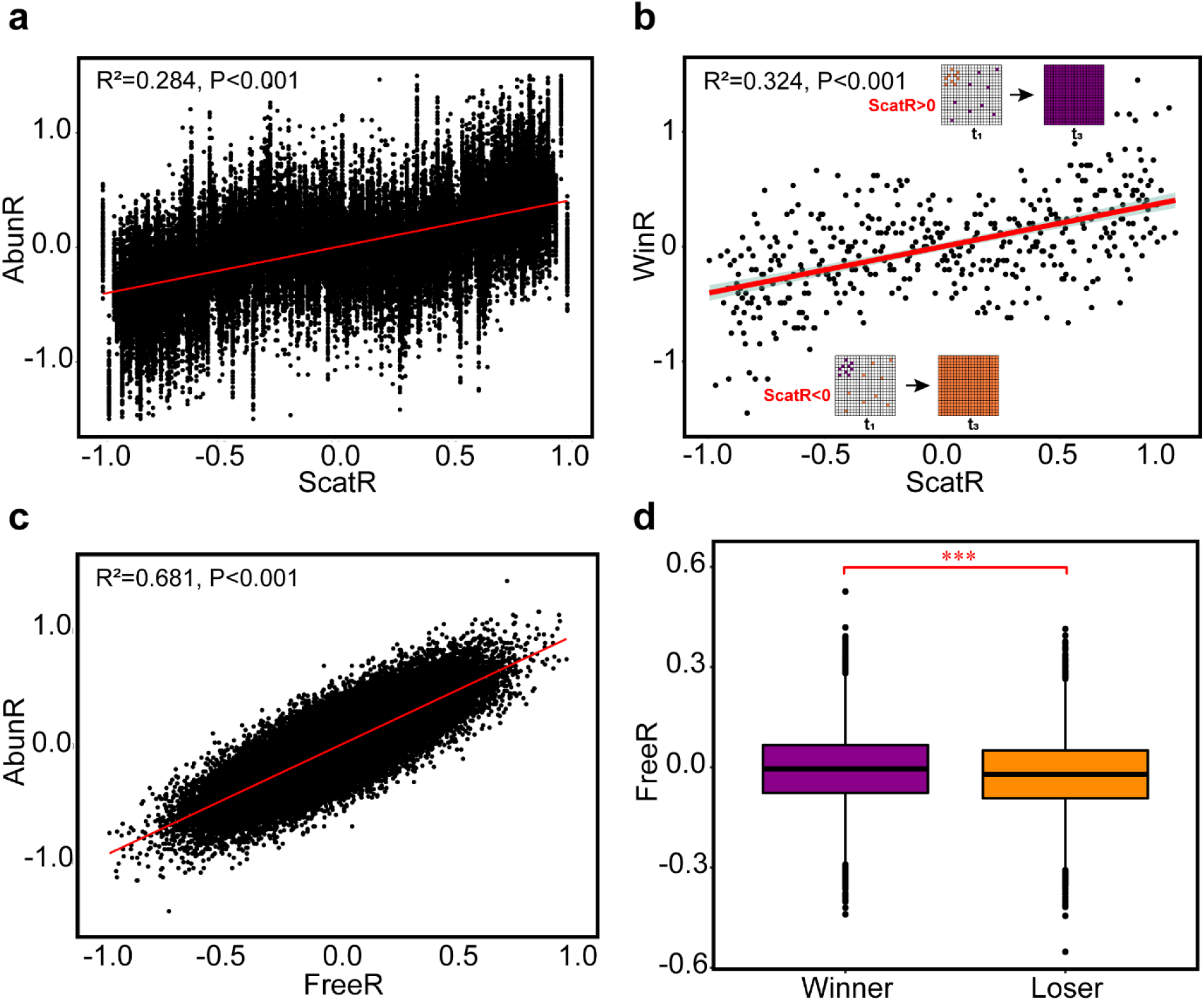
Effect of the initial cell distribution and expanding direction of the daughter cells on competitive success of one population. **a.** Correlation between ScatR and AbunR. Results represented the sum of 21500 simulations containing 215 different initial cell distribution, of which the ScaR values varied from −1.053 to 1.053. **b.** Correlation between ScatR and WinR. Data were generated from simulations identical to those shown in a., and each WinR value was summarized from the competition outcome of 100 replicated simulations with a given initial cell distribution. **c.** Correlation between FreeR and AbunR. Results represented the sum of 36300 independent simulations with 363 different initial cell distribution, but their ScaR values were all equal to zero (Fig. S3). **d.** Comparison between ‘expansion freedom’ of the winning population and that of the losing population. Competition outcomes were generated from simulations identical to those shown in **c**. Statistical analysis was performed using a two-sample Student’s t-test: ***, p < 0.001.

#### Higher degree of expansion freedom helps populations to win space competition

In addition to initial distance, we found that for a given initial distribution, AbunR considerable varied across different expansion processes, indicating that in addition to the randomness in the initial cell distribution, the random events occurring during population expansion in the “occupation stage” also affected the competition outcome. Our model assumed that after a successful division of one cell, all eight grid boxes around the mother cell are randomly selected to accommodate the newly born cell (Fig. 1a; assumption 2). If the selected box has been occupied, the newborn cell will compete for the box with the aborigine of the box and has a 50% probability to survive (assumption 3). We defined the number of empty boxes surrounding the newborn cell as the degree of expansion freedom. Thus, if the daughter cell possesses a higher degree of expansion freedom, the probability for its offspring to survive will be higher (Fig. 1d; the purple cell). In contrast, if the degree of expansion freedom of the daughter cell is low, it has to compete for space with other cells for further reproduction and expansion, which should be less favored for the space competition afterwards (Fig. 1d; the orange cell). This assumption leads to a prediction that the population whose daughter cells possess a higher degree of expansion freedom will be more likely to win the competition. To test this prediction, we defined the expansion freedom asymmetry index, FreeR, of a population as the difference in expansion freedom between a population and its competitor.

We selected 363 initial cell distributions with a zero ScatR (Fig. S3) from 1000000 random distributions and performed 100 replicated simulations with each initial distribution. During these simulations, we recorded the degrees of expansion freedom of every newborn cell during the ‘occupation stage’ (Fig. 1d; Fig. S4) and then compared the FreeR of a population with its AbunR in each simulation. We observed a strong positive relationship between FreeR and AbunR (Fig. 2c; R^2^=0.681, P<0.001), suggesting that the population with greater ‘expansion freedom’ will occupy more space at t2. Furthermore, the winning populations exhibited a significantly higher FreeR than the populations losing the competitions (Fig. 2d; t-value=18.406, df=53074, P<0.001), which is consistent with our prediction.

Together, these results demonstrated that the randomness during the “occupation stage” of spatial competition, including the initial scatter level and the degree of expansion freedom, can affect the outcome of competition for space.

#### Populations with higher ‘Space Accessibility’ have higher winning probability in spatial competition

Because both the initial scatter level and the degree of expansion freedom affect the number of free boxes that surrounded the populations at each time point, we then searched for a more general parameter that considers both of the two factors. We applied a mathematical induction algorithm to define a new parameter, Space Accessibility (SA_k,t_, Fig. S5). Individuals of a population are further away from the aggregation area, resulting in a higher ‘Space Accessibility’. The ‘Space Accessibility’ at each time point assesses the probability of the cells of one population colonizing all the empty space in the subsequent competition from this time point, which reflects the ease with which daughter cells move into these empty positions. Next, we integrated SA_k,t_ value over time (obtaining SA) for each population. To estimate which population generally was more likely to occupy the empty positions during the “occupation stage”, we next defined an index called SAR. A SAR index greater than zero indicates that a population has a higher probability of reaching empty positions than its competitor across the “occupation stage”.

To investigate whether the difference in ‘Space Accessibility’ affects the competition outcome, we performed 20000 simulations covering SAR values ranging from −1.754 to 1.754. In these simulations, we found that ScatR (Fig. S6a; R^2^=0.283, P<0.001), as well as FreeR (Fig. S6b; R^2^=0.986, P<0.001), is positively correlated with the SAR, suggesting that the values of SAR reflect the values of both ScatR and FreeR. To test the influence of ‘Space Accessibility’ for competition, we next analyzed the relationship between SAR of a population and its AbunR at t_2_ time point. The results showed an extremely significant positive correlation between the SAR and AbunR (Fig. 3a; R^2^=0.830, P<0.001), suggesting that the population with higher ‘Space Accessibility’ will occupy more space at t_2_. The correlation coefficient between AbunR and SAR was higher than the coefficients of both AbunR-ScatR and AbunR-FreeR, indicating that SAR represents a more suitable parameter to evaluate competition outcomes. Furthermore, the winning populations possessed a significantly higher SAR than the populations losing the competitions in all simulations, (Fig. 3b; t-value=40.256, df=39998, P<0.001), further indicating that the ‘Space Accessibility’ predicted the outcome of spatial competition between two populations with a high degree of reliability.

**Fig. 3.**
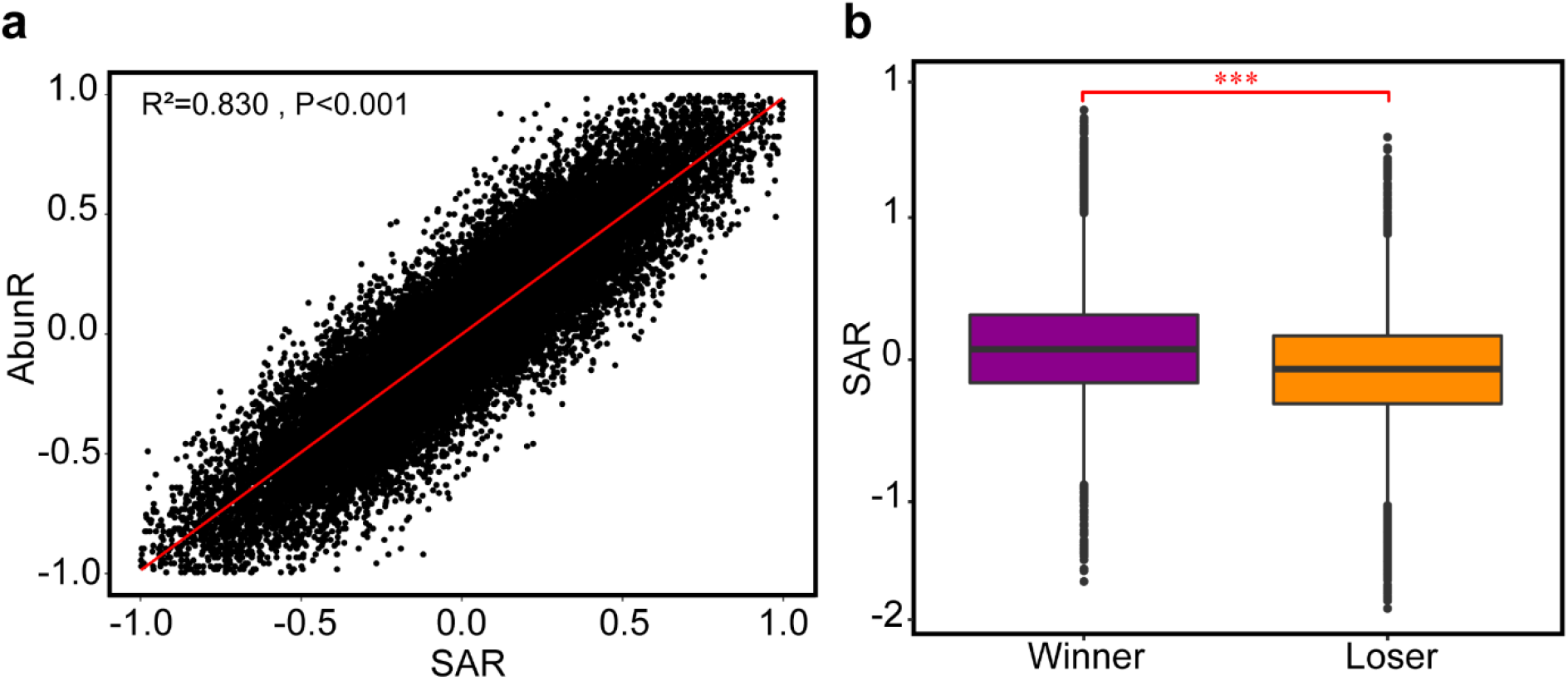
Effect of the ‘Space Accessibility’ on the competitive success of a population. **a.** Correlation between SAR and AbunR. Results represented the sum of 20000 independent simulations. **b**. Comparison between the SAR of the winning population and that of the failed population. Competition outcomes were generated from simulations identical to those shown in a. Statistical analysis was performed using a two-sample Student’s t-test: ***, p < 0.001.

We also performed numerous well-designed simulations (See Supplementary Information S2 for details) to test whether the effect of ‘Space Accessibility’ on the outcome of spatial competition is statistically significant under various initial conditions (robustness test), including varied initial growth rate, total number of initial cells, as well as the size of the space (Table S1). Our analysis showed that the effect of ‘Space Accessibility’ on the outcome of spatial competition is significant (Table S1; Fig. S7), and largely unperturbed by changes in the initial growth rate and space size and the initial total number of cells (not exceed to 10% of the maximum population size). In summary, colonizing space in an anti-aggregation manner contributes to microbial competitive success.

#### A ‘smart population’ occupies more space

Next, we tested whether ‘Space Accessibility’ determines the competition outcome from another perspective. We designed simulations of competition between SmartBac and NormalBac (See Materials and methods for more details). We hypothesized that SmartBac would obtain a higher competitive edge from its superior strategy of space colonization, and win the competition for space against NormalBac.

As expected, SmartBac won 10227 times during 14008 mathematical simulations, accounting for 73.01%. In these simulations, the SAR values of the SmartBac were significantly higher than those of the NormalBac (Fig. 4a; t-value=304.471, df=28014, P<0.001). Furthermore, the AbunR values of SmartBac were also significantly higher than those of NormalBac (Fig. 4b; t-value=383.574, df=28014, P < 0.001). Together, the results further indicated that microbial colonization of space in an anti-aggregation manner helps winning the competition.

**Fig. 4.**
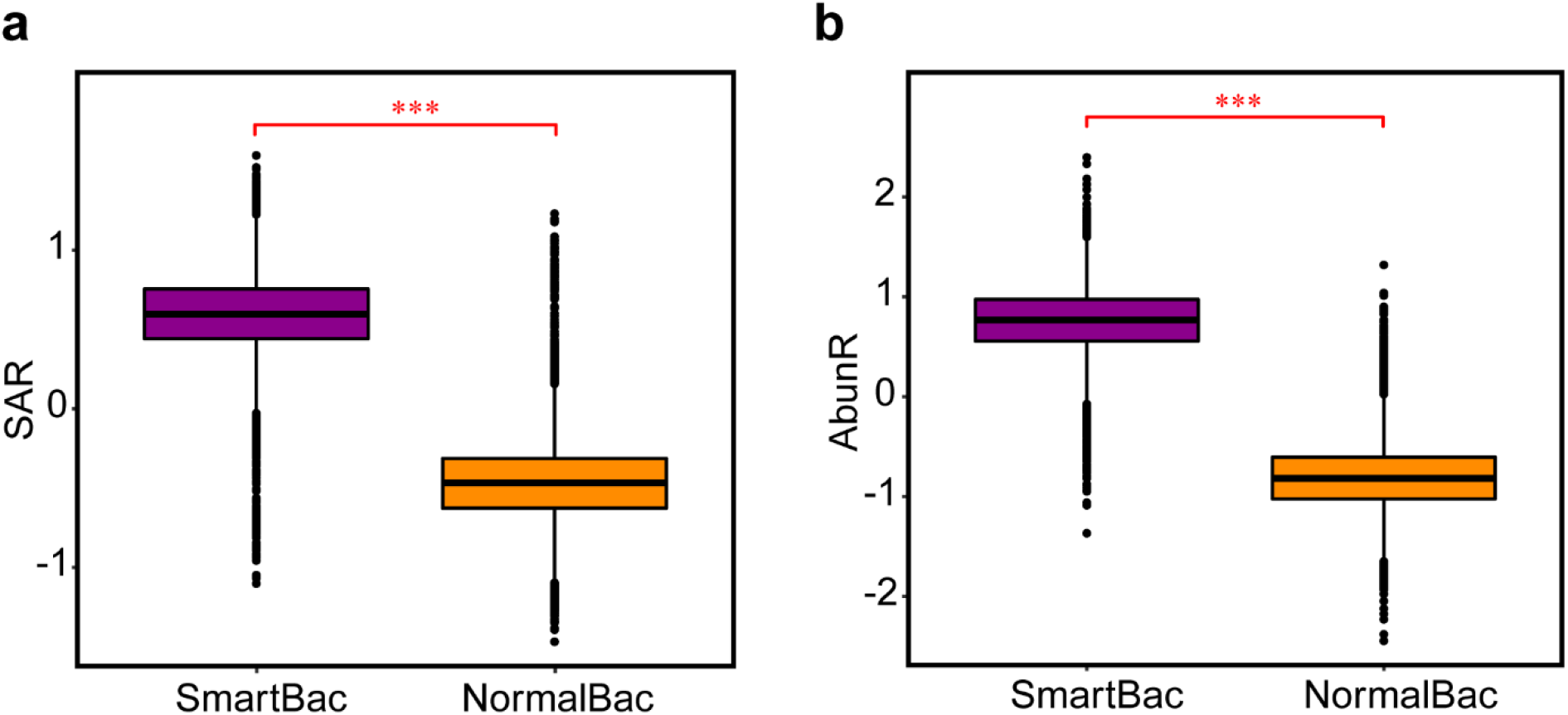
Microbial colonization of space in an anti-aggregation manner contributes to the competitive success of the SmartBac against the NormalBac. **a.** Comparison of the SAR of the SmartBac and NormalBac in the “occupation stage”. **b.** Comparison of the AbunR at t_2_ of the SmartBac and NormalBac. Results represented the sum of 14008 independent simulations and statistical analysis was performed by two-sample Student’s t-test: ***, P < 0.001.

### Space colonization manners, growth rates and initial abundances synergistically affect spatial competition

It is well established that microbial competition for space is influenced by the growth rate and initial abundance of the competing populations. The population possessing a faster growth rate, or higher initial abundance will outcompete other strains present within the newly occupied space. To assess the relative contribution of space colonization manners, different growth rates and initial abundances in spatial competition, to microbial competitive success, we performed simulations in which the growth rates and initial abundances of the two populations were set to be different. In these simulations, we defined the parameter GroR as the difference in growth rate between a population and its competitor, as well as defined InifR to characterize the difference in initial abundances (Fig. 5a). In addition, we calculated the SAR value of each competitive population in each simulation to quantify the asymmetry of ‘Space Accessibility’.

**Fig. 5.**
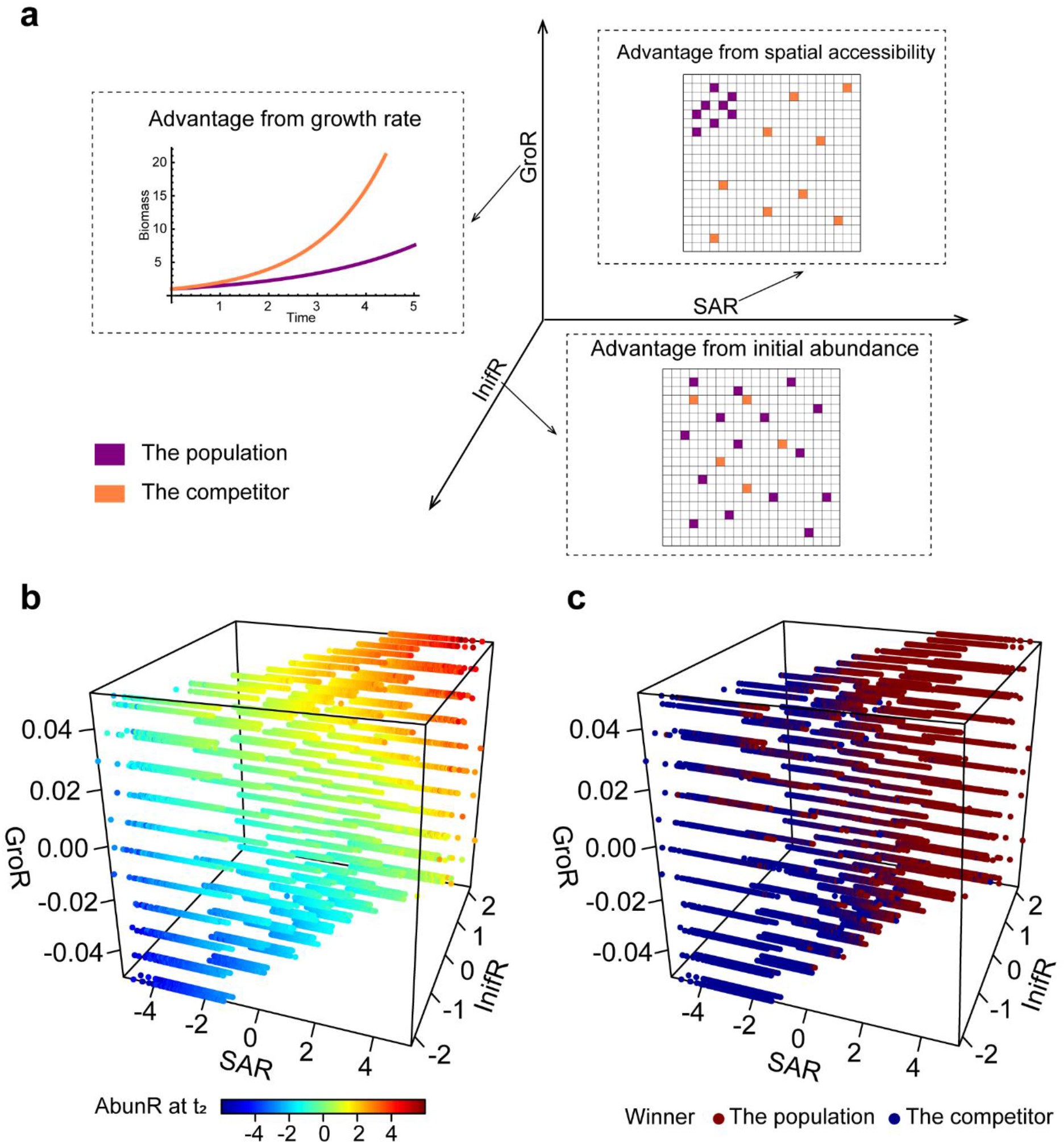
Comparison of the effect of space colonization manners with the effect of the varied growth rate and initial abundance. **a.** Diagram indicating the meaning of the defined three parameters. Values of each parameter greater than 0 denote that the focal population possesses the corresponding competitive edge compared with its competitor. **b-c.** Comparison of the relative importance of the space colonization manners, the growth rate and higher initial abundance for the outcome of microbial competition. Gradients of SAR, GroR, as well as InifR were set in 89100 simulations and each point indicated the simulation result in the corresponding set of the three parameters. Values of AbunR (**b**), as well as final competition outcomes (**c**) were also recorded to estimate how these three factors collectively affect the microbial competition.

As shown in Fig. 5, even when one population exhibited lower growth rate, or was characterized by lower initial abundance, colonization of space in an anti-aggregation manner, such as choosing positions for new cells to have a higher ‘Space Accessibility’, would neutralize these disadvantages and allow this population to occupy more space at t_2_ (Fig. 5b), thus winning the spatial competition (Fig. 5c). The collinearity analysis showed that when strains differed in their initial abundances, SAR and InifR exhibited significant collinearity (VIF=13.062).

To eliminate this collinearity effect, we generalized our definition of SAR by defining a new parameter perSAR (See Supplementary Information S1 for details), which is equal to SAR when the initial cell number of both populations (InifR = 0), but allows for better quantification of the asymmetry of ‘Space Accessibility’ when InifR is unequal to zero. A subsequent collinearity test showed that the collinearity among the variables perSAR, GroR and InifR disappeared (Table S2). Moreover, the population possessing a higher perSAR value was characterized by a higher probability for ultimate survival at the end of the simulation, and won the competition even when its growth rate or initial abundance was lower (Fig. S8).

We next performed multiple regression analysis to quantify the relative contributions of the three factors during spatial competition (Table S2). Our analysis showed that the ratio of relative contributions of perSAR, GroR and InifR to AbunR were approximately 1.027, 55.393 and 1.027 (1:53.94:1), suggesting that the competitive disadvantage derived from lower GroR of a population can be eliminated by possessing 53.94 times higher perSAR, and the competitive disadvantage derived from lower InifR can be neutralized by 1-time higher perSAR. Together, these results indicated that microbial colonization of space in an anti-aggregation manner can benefit the competitive success of slow-growing species or species possessing lower seeding abundance.

In summary, compared with the evident competitive edge derived from a faster growth rate and higher initial abundance of one competitor, colonization of space in an anti-aggregation manner (e.g. possessing higher ‘Space Accessibility’) also play a critical role in determining the success rate during competition for space between microbial strains.

## Discussion

In this study, we investigated whether and how space colonization manners affect the outcome of spatial competition among different microbial populations, which is ecologically significant to predict the dynamics of natural communities. Our results suggested that smart populations, which colonize space in an anti-aggregation manner, can win the spatial competition even if they grow slightly slower than their competitors.

We considered exploring the spatially related abiotic factors of spatial competition, which were inspired by a traditional Chinese board game, *Go*. In the game *Go*, two players need to rationally consider the strategy of how to place their game pieces in order (strategically determines where to place a piece at each step(Baker 2008)) to win the game by occupying more area on the board. This process is in close analogy to the ecological processes that two biological populations competitively colonize an uninhabited space. Inspired by the golden rule(Baker 2008) of winning a *Go* game, “golden corner, silver side and grass belly”, we proposed the hypothesis that manners of organisms colonizing free space, such as choosing initial positions and subsequent directions of expansion, play a significant role in their competitive outcome.

Initial spatial pattern and random processes during microbial population expansion are important for the spatial competition(O *et al.* 2007; Lloyd & Allen 2015). For example, one recent study explored how an *Escherichia coli* population colonized the surface of a flat agarose pad and investigated how two *E. coli* populations compete for limited space(Lloyd & Allen 2015). In that study, cells with smaller initial patches are more likely to be winners, which agrees with our model prediction, as more scattered initial distribution normally leads to smaller patches at the beginning of colonization. In addition, random processes such as spatial wandering of so-called ‘pioneers’ at the expanding frontier of a colony(O *et al.* 2007; Chu *et al.* 2019), will determine the spatial competition among the founder cells. Our findings presented here suggest that the direction towards which the newly divided cells migrate represents an important random event in the expansion of a colony, critically affecting the spatial competition between different populations. The populations generating offsprings with greater ‘expansion freedom’ will occupy more space at t_2_ and thus gaining an advantage over other strains competing for space. Therefore, space colonization is a vital stochastic factor that governs the interactions between competing microbes, as well as the structure of their communities.

Ecological competition can result in the evolution of phenotypes. Several studies using experimental evolution have documented evolution in spatial competition. For example, a mutant repeatedly arose in a biofilm formed by *Pseudomonas fluorescens* Pf0-1, able to maintain a presence at the surface of the biofilm, thus gaining access to limiting nutrients and space(Kim *et al.* 2014). Here, we hypothesized that the evolution of colonizing space in an anti-aggregation manner may benefit the spatial competition of microbes. Our simulations indicated that an evolved population (SmartBac), that always migrated to the grids with higher ‘Space Accessibility’, will be selectively favored (Fig. 4). Recent studies have provided clues supporting this evolution strategy. Quorum sensing (QS), a signaling system that regulates gene expression and coordinates population behavior in response to changes in population density, is very common among microbes(Schuster *et al.* 2013; Papenfort & Bassler 2016; Abisado *et al.* 2018; Mukherjee & Bassler 2019). QS signals can be used to direct the free areas, that the free area is larger, the concentration of QS signals should be lower. Exopolysaccharides (EPS), acting as ‘molecular glue’, promote surface attachment and affect surface exploration(Ma *et al.* 2007; Byrd *et al.* 2009; Byrd *et al.* 2010). One study combining the cell-tracking technique and computer simulations showed that *P. aeruginosa* deposits a trail of Psl as it moves on a surface, which influences the surface motility of subsequent cells that encounter these trails(Zhao *et al.* 2013). The slime mold *Physarum polycephalum* can navigate complex labyrinths to find the optimal path to a food source(Alim *et al.* 2017). In these cases, microorganisms can sense the population density and migrate in a directional manner, suggesting that the possibility of the evolution for colonizing space in an anti-aggregation manner. To test this hypothesis, a long-term experimental evolution assay performing spatial competition of two populations at the individual level should be designed.

Although we built our model based on a set of assumptions considering the lifestyle of microorganism, specific rules derived from the simulations also have implications for understanding space colonization of macroorganism. Data of 54 natural forest areas from ForestGEO (https://forestgeo.si.edu/) confirms that seeding with more scattered initial distribution contributes to faster space colonization of trees, which is consistent with the conclusion of our microbial competition model (Fig. S9). Therefore, our findings can also be generalized to explain how multicellular individuals compete for space and may help to design ecological restoration strategies, such as artificial forestation.

Competition among organisms has a large impact on species diversity(Maynard *et al.* 2017). Traditionally, competitive edges were mostly attributed to faster growth rates(Sebens 1982). However, how slower-growing species survive in nature has remained an unsolved conundrum. Our analysis of spatial competition of two populations indicated that the competitive disadvantage derived from a slower growth rate can be neutralized by higher ‘Space Accessibility’. As a result, colonizing space in an anti-aggregation manner will benefit the competitive success of a slower-growing species. This result provided a novel perspective that the randomness in space colonization, or evolving a smarter manner for space colonization, may contribute to the survival of those slow-growing species. Spatial-structured environments, such as biofilm or soil, commonly exhibit higher spatial heterogeneity(Dzubakova *et al.* 2018; CJ *et al.* 2019; Ye *et al.* 2019; Jiang *et al.* 2020), characterized by numerous homogeneous microhabitats(Rybicki *et al.* 2020). The competition outcome in each microhabitat varies due to random space colonization, allowing the co-existence of species with different growth rates at a large spatial scale. Therefore, our results also provide novel insights into the maintenance of biodiversity in spatial-structured environments.

Our results clearly demonstrate that disadvantaged strains use innovative strategies when colonizing newly discovered spaces, compensating for disadvantageous biotic conditions, and thus considerably improving its changes in the evolutionary arms race. These findings shed light on the role of spatial positioning in maintaining biodiversity within natural communities, as well as provides a new insight on how spatial competition between different populations drives evolutionary innovation.

## Acknowledgements

This work was supported by National Key R&D Program of China (2018YFA0902100 and 2018YFA0902103), National Natural Science Foundation of China (91951204, 31770120, 31770118 and 31761133006) and High-performance Computing Platform of Peking University.

## Author contributions

M.W. conceived the model, and X.L. designed and carried it out. X.L. collected and analyzed the data. X.L., M.W. and Y.N. co-wrote the paper, and X.L.W. revised it. All authors discussed the results and commented on the paper.

## References

1. Abisado, R.G., Benomar, S., Klaus, J.R., Dandekar, A.A. & Chandler, J.R. (2018). Bacterial Quorum Sensing and Microbial Community Interactions. mBio, 9.

2. Abreu, C.I., Friedman, J., Andersen Woltz, V.L. & Gore, J. (2019). Mortality causes universal changes in microbial community composition. Nat Commun, 10, 2120.

3. Ali, S.S. & Vidhale, N. (2013). Bacterial Siderophore and their Application: A review Int J Curr Microbiol App Sci 2, 303–312.

4. Alim, K., Andrew, N., Pringle, A. & Brenner, M.P. (2017). Mechanism of signal propagation in Physarum polycephalum. Proc Natl Acad Sci U S A, 114, 5136–5141.

5. Baalen, M.V. & Yamauchi, A. (2019). Competition for resources may reinforce the evolution of altruism in spatially structured populations. Math Biosci Eng, 16, 3694–3717.

6. Baker, K. (2008). The Way to Go: How to Play the Asian Game of Go. Seventh Edition edn. NY: American Go Association, New York.

7. Benayahu, Y. & Loya, Y. (1981). Competition for Space among Coral-Reef Sessile Organisms at Eilat, Red Sea. Bulletin of Marine Science, 31, 514–522.

8. Bernhardt, J.R., Kratina, P., Pereira, A.L., Tamminen, M., Thomas, M.K. & Narwani, A. (2020). The evolution of competitive ability for essential resources. Philos Trans R Soc Lond B Biol Sci, 375, 20190247.

9. Blount, Z.D., Barrick, J.E., Davidson, C.J. & Lenski, R.E. (2012). Genomic analysis of a key innovation in an experimental Escherichia coli population. Nature, 489, 513–518.

10. Burson, A., Stomp, M., Greenwell, E., Grosse, J. & Huisman, J. (2018). Competition for nutrients and light: testing advances in resource competition with a natural phytoplankton community. Ecology, 99, 1108–1118.

11. Byrd, M.S., Pang, B., Mishra, M., Swords, W.E. & Wozniak, D.J. (2010). The Pseudomonas aeruginosa exopolysaccharide Psl facilitates surface adherence and NF-kappaB activation in A549 cells. mBio, 1.

12. Byrd, M.S., Sadovskaya, I., Vinogradov, E., Lu, H., Sprinkle, A.B., Richardson, S.H. et al. (2009). Genetic and biochemical analyses of the Pseudomonas aeruginosa Psl exopolysaccharide reveal overlapping roles for polysaccharide synthesis enzymes in Psl and LPS production. Mol Microbiol, 73, 622–638.

13. Campbell, B.J., Yu, L., Heidelberg, J.F. & Kirchman, D.L. (2011). Activity of abundant and rare bacteria in a coastal ocean. Proc Natl Acad Sci U S A, 108, 12776–12781.

14. Chu, S., Kardar, M., Nelson, D.R. & Beller, D.A. (2019). Evolution in range expansions with competition at rough boundaries. J Theor Biol, 478, 153–160.

15. CJ, B., AR, E., SH, W., SA, R., JR, B., S, D. et al. (2019). Respiratory Heterogeneity Shapes Biofilm Formation and Host Colonization in Uropathogenic Escherichia coli. mBio, 10, e02400–02418.

16. Crowley, P.H., Davis, H.M., Ensminger, A.L., Fuselier, L.C., Kasi Jackson, J. & Nicholas McLetchie, D. (2004). A general model of local competition for space. Ecology Letters, 8, 176–188.

17. Davis, K.E., Sangwan, P. & Janssen, P.H. (2011). Acidobacteria, Rubrobacteridae and Chloroflexi are abundant among very slow-growing and mini-colony-forming soil bacteria. Environ Microbiol, 13, 798–805.

18. Denboer, P.J. (1986). The present status of the competitive exclusion principle. Trends in Ecology & Evolution, 1, 25–28.

19. DK, B. (1998). Nutrient Uptake by Microorganisms according to Kinetic Parameters from Theory as Related to Cytoarchitecture. Microbiol Mol Biol Rev, 62, 636–645.

20. Dzubakova, K., Peter, H., Bertuzzo, E., Juez, C., Franca, M.J., Rinaldo, A. et al. (2018). Environmental heterogeneity promotes spatial resilience of phototrophic biofilms in streambeds. Biol Lett, 14.

21. Eickhoff, M.J. & Bassler, B.L. (2020). Vibrio fischeri siderophore production drives competitive exclusion during dual-species growth. Mol Microbiol, 114, 244–261.

22. Gause, G.F. (1932). Experimental studies on the struggle for existence I Mixed population of two species of yeast. Journal of Experimental Biology, 9, 389–402.

23. Henson, M.W., Lanclos, V.C., Faircloth, B.C. & Thrash, J.C. (2018). Cultivation and genomics of the first freshwater SAR11 (LD12) isolate. ISME J, 12, 1846–1860.

24. Jiang, Y., Zhang, B., Wang, W., Li, B., Wu, Z. & Chu, C. (2020). Topography and plant community structure contribute to spatial heterogeneity of soil respiration in a subtropical forest. Sci Total Environ, 733, 139287.

25. Kim, W., Racimo, F., Schluter, J., Levy, S.B. & Foster, K.R. (2014). Importance of positioning for microbial evolution. Proc Natl Acad Sci U S A, 111, E1639–1647.

26. Kreft, J.-U., Booth, G. & Wimpenny, J.W.T. (1998). BacSim, a simulator for individual-based modelling of bacterial colony growth. Microbiology, 144, 3275–3287.

27. Lax, S., Abreu, C.I. & Gore, J. (2020). Higher temperatures generically favour slower-growing bacterial species in multispecies communities. Nat Ecol Evol, 4, 560–567.

28. Lirman, D. (2001). Competition between macroalgae and corals: effects of herbivore exclusion and increased algal biomass on coral survivorship and growth. Coral Reefs, 19, 392–399.

29. Lloyd, D.P. & Allen, R.J. (2015). Competition for space during bacterial colonization of a surface. J R Soc Interface, 12, 0608.

30. Ma, L., Lu, H., Sprinkle, A., Parsek, M.R. & Wozniak, D.J. (2007). Pseudomonas aeruginosa Psl is a galactose- and mannose-rich exopolysaccharide. J Bacteriol, 189, 8353–8356.

31. MacLean, R.C. & Gudelj, I. (2006). Resource competition and social conflict in experimental populations of yeast. Nature, 441, 498–501.

32. Maynard, D.S., Bradford, M.A., Lindner, D.L., van Diepen, L.T.A., Frey, S.D., Glaeser, J.A. et al. (2017). Diversity begets diversity in competition for space. Nat Ecol Evol, 1, 156.

33. Mukherjee, S. & Bassler, B.L. (2019). Bacterial quorum sensing in complex and dynamically changing environments. Nat Rev Microbiol, 17, 371–382.

34. Niehus, R., Picot, A., Oliveira, N.M., Mitri, S. & Foster, K.R. (2017). The evolution of siderophore production as a competitive trait. Evolution, 71, 1443–1455.

35. O, H., P, H., S, R. & Dr, N. (2007). Genetic drift at expanding frontiers promotes gene segregation. Proc Natl Acad Sci U S A, 104, 19926–19930.

36. Oliveira, N.M., Martinez-Garcia, E., Xavier, J., Durham, W.M., Kolter, R., Kim, W. et al. (2015). Biofilm Formation As a Response to Ecological Competition. PLoS Biol, 13, e1002191.

37. P, A. & K, B. (1981). Competitive Advantage Provided by Bacterial Motility in the Formation of Nodules by Rhizobium meliloti. J Bacteriol, 148, 728–729.

38. Papenfort, K. & Bassler, B.L. (2016). Quorum sensing signal-response systems in Gram-negative bacteria. Nat Rev Microbiol, 14, 576–588.

39. Paquette, S.J., Zaheer, R., Stanford, K., Thomas, J. & Reuter, T. (2018). Competition among Escherichia coli Strains for Space and Resources. Vet Sci, 5.

40. Petrova, O.E. & Sauer, K. (2012). Sticky situations: key components that control bacterial surface attachment. J Bacteriol, 194, 2413–2425.

41. Ratzke, C. & Gore, J. (2018). Modifying and reacting to the environmental pH can drive bacterial interactions. PLoS Biol, 16, e2004248.

42. RJ, S. & DJ, M. (2015). Limiting resources in sessile systems: food enhances diversity and growth of suspension feeders despite available space. Ecology, 26, 819–827.

43. Rybicki, J., Abrego, N. & Ovaskainen, O. (2020). Habitat fragmentation and species diversity in competitive communities. Ecol Lett, 23, 506–517.

44. Scharf, B.E., Hynes, M.F. & Alexandre, G.M. (2016). Chemotaxis signaling systems in model beneficial plant-bacteria associations. Plant Mol Biol, 90, 549–559.

45. Schluter, J., Nadell, C.D., Bassler, B.L. & Foster, K.R. (2015). Adhesion as a weapon in microbial competition. ISME J, 9, 139–149.

46. Schuster, M., Sexton, D.J., Diggle, S.P. & Greenberg, E.P. (2013). Acyl-homoserine lactone quorum sensing: from evolution to application. Annu Rev Microbiol, 67, 43–63.

47. Sebens, K.P. (1982). Competition for Space: Growth Rate, Reproductive Output, and Escape in Size. The American Naturalist, 120, 189–197.

48. Si, C., Zhang, L.-M. & Yu, F.-H. (2019). Effects of physical space and nutrients on the growth and intraspecific competition of a floating fern. Aquatic Ecology, 53, 295–302.

49. Smith, G.R., Steidinger, B.S., Bruns, T.D. & Peay, K.G. (2018). Competition-colonization tradeoffs structure fungal diversity. ISME J, 12, 1758–1767.

50. Wang, M., Geng, S., Hu, B., Nie, Y. & Wu, X.L. (2021a). Sessile bacterium unlocks ability of surface motility through mutualistic interspecies interaction. Environ Microbiol Rep, 13, 112–118.

51. Wang, M., Liu, X., Nie, Y. & Wu, X.L. (2020). Selfishness driving reductive evolution shapes interdependent patterns in spatially structured microbial communities. ISME J.

52. Wang, M., Nie, Y. & Wu, X.L. (2021b). Extracellular heme recycling and sharing across species by novel mycomembrane vesicles of a Gram-positive bacterium. ISME J, 15, 605–617.

53. Wang, M., Nie, Y. & Wu, X.L. (2021c). Membrane vesicles from a Dietzia bacterium containing multiple cargoes and their roles in iron delivery. Environ Microbiol, 23, 1009–1019.

54. Ye, H., Lu, C. & Lin, Q. (2019). Investigation of the spatial heterogeneity of soil microbial biomass carbon and nitrogen under long-term fertilizations in fluvo-aquic soil. PLoS One, 14, e0209635.

55. Zhao, K., Tseng, B.S., Beckerman, B., Jin, F., Gibiansky, M.L., Harrison, J.J. et al. (2013). Psl trails guide exploration and microcolony formation in Pseudomonas aeruginosa biofilms. Nature, 497, 388–391.

