## Supplementary Information for "Successful microbial colonization of space using an anti-aggregation strategy"

### for

### S1 Simulations designed for investigating the effect of different colonization manners on spatial competition

#### *Simulations for investigating the effect of initial cell distribution on spatial competition*

Firstly, a non-dimensional parameter, ScatR, was defined to assess the asymmetry of the scatter level of the initial cell distribution between a population and its competitor, calculated by:

$$a_1 = \sqrt{\frac{\sum_{i=1}^{n_1}(x_{1i} - \overline{x_1})^2 + (y_{1i} - \overline{y_1})^2}{n_1}}$$

$$a_2 = \sqrt{\frac{\sum_{i=1}^{n_2}(x_{2i} - \overline{x_2})^2 + (y_{2i} - \overline{y_2})^2}{n_2}}$$

$$ScatR = \log(\frac{a_1}{a_2})$$

In which, $(x_{1i}, y_{1i})$ represents the position coordinate of the $i$th individual of the focus population, while $(x_{2i}, y_{2i})$ represents that of its competitor. $n_1$ and $n_2$ are the initial cell numbers of the two populations, respectively. $a_1$ and $a_2$ are the average Euclidean distance between cells of two populations respectively, which characterize the scattered level of initial cell distribution of each population. Therefore, ScatR reflects the scatter asymmetry of one population in the initial distribution. The ScatR greater than 0 indicates that the population is initially distributed more scatteredly than its competitor, and the absolute value of ScatR represents the degree of the difference in the scatter level of the initial cell distribution between the two populations.

Secondly, to obtain initial cell distributions with different relative scattered level, 1000000 cell distributions were randomly generated and ScatR values of these distributions were calculated (Fig. S3). According to these calculations, 215 distributions were selected, which covers a gradient of ScatR values ranged from -1.053 to 1.053 (in other words, when ScatR increased by 0.01, approximately one initial distribution was selected).

Thirdly, 100 replicated simulations were performed initialized with each cell distribution and 21500 simulations in total were run. AbunR and WinR values of these simulations then were calculated and the linear correlation between ScatR, AbunR and

WinR, were analyzed.

***Simulations for investigating the effect of 'expansion freedom' on spatial competition***

In order to investigate the effect of degree of 'expansion freedom' on the competition outcome, 363 cell distributions were selected from the 1000000 ones generated from the above part, of which the ScatR values were all equal to zero (Red line in Fig. S3). Also, 100 replicated simulations were performed, and 36300 simulations were performed initialized with these cell distributions in total. During simulations, the moving direction (position coordinates) of every newly born cells were tracked in detail. After simulations, a parameter FreeR was defined to characterize the difference in the degree of 'expansion freedom' between one population and its competitor, given by

$$\mathrm{FreeR} = \log\left(\frac{\sum_{t=0}^{t_2} \overline{freedom_{1,t}}}{\sum_{t=0}^{t_2} \overline{freedom_{2,t}}}\right)$$

Here, the $\overline{freedom_{1,t}}$ is the average number of empty grids around the daughter cells born in time point $t$ of the focal population, while $\overline{freedom_{2,t}}$ is that number of its competitor. A summation of the $\overline{freedom}$ values across the "occupation stage" reflects average empty-position numbers surrounding the population during the spatial competition. Here, the FreeR index greater than zero indicates that the population possesses greater 'expansion freedom' against its competitor in the given simulation, and the higher absolute value suggests a higher difference in expansion freedom between the two populations. Based on this definition, FreeR evaluates the competitive edge derived from the asymmetric 'expansion freedom' of one population against its competitor across the "occupation stage".

Finally, linear correlation between AbunR and FreeR of each simulation, as well as

FreeR values between the competition outcome of those winning population and losing population, was statistically analyzed.

***Simulations for investigating the effect of 'Space Accessibility' on spatial competition***

In order to integrate the effect of initial cell distribution and 'expansion freedom', a new parameter, named 'Space Accessibility', was defined. Firstly, $SA_{k,j,t}$ was defined as the maximum probability that the offspring cell of the *j*th individual of the *k*th population occupies the all unoccupied grids at time point *t*. To calculate$SA_{k,j,t}$, the all unoccupied grids was divided into 19 layers centered with the grid of the *j*th individual (Since the whole space was a 20×20 array; Fig. S5). As shown in Fig. S5, the maximum probability of the offspring of the *j*th individual to occupy a grid in the *i*th layer, $P_{i,j}$, was calculated by

$$P_{ij} = \frac{\sum_{m=0}^{i-1} 3^m}{i} \times (\frac{1}{8})^i$$

Next, $SA_{k,j,t}$ was derived by

$$SA_{k,j,t} = \sum_{i=1}^{19} P_{i,j} \times G_{k,i,j,t}$$

Here, $G_{k,i,j,t}$ is the number of the empty grids in the *i*th layer surrounding the *j*th individual of the *k*th population at time point *t*. To assess the maximum probability of the cells of the *k*th population occupy all the empty space at time point *t* in the follow-up steps, 'Space Accessibility' of the *k*th population was then defined as the summation of the $SA_{k,j,t}$ value of every individuals of the *k*th population at time point *t*, given by

$$SA_{k,t} = \sum_{j=1}^{n_{k,t}} SA_{k,j,t}$$

Here, $n_{k,t}$ is the number of individuals of the *k*th population at time point *t*. Finally, the asymmetry of 'Space Accessibility', SAR, was defined to collectively evaluated the competitive edge derived from 'Space Accessibility' of the population across whole "occupation stage", given by

$$\mathrm{SAR} = \log\left(\frac{\sum_{t=0}^{t_2} SA_{1,t}}{\sum_{t=0}^{t_2} SA_{2,t}}\right)$$

Here, $SA_{1,t}$ and $SA_{2,t}$ are the 'Space Accessibility' for the focal population and its competitor at time point t, respectively. By this definition, SAR greater than 0 means that the population generally possess higher 'Space Accessibility' than its competitor. To investigate how the 'Space Accessibility' affect the outcome of spatial competition between two populations, 200 initial cell distributions were selected, which covers a gradient of ScatR values ranged from -1.0 to 1.0. 100 replicated simulations were performed initialized with each cell distribution and 20000 simulations in total were run. In each simulation step, $SA_{1,t}$ and $SA_{2,t}$ values were calculated by the above methods, implemented by custom C++ code (https://github.com/Neina-0830/BacGo-model). After simulations, SAR values, AbunR, as well as WinR values of these simulations were calculated and analyzed.

***Simulations for comparing the effect of space colonization manners, growth rates, and initial frequencies on spatial competition***

To compare the relative importance of the effect of space colonization manners, different growth rates and different initial abundances for microbial competition, three parameters were defined to assess the different asymmetries of the two populations in the above three aspects. Specifically, GroR was defined to assess the asymmetry of growth rate of each population, given by

$$\mathrm{GroR} = \log(\frac{Gro_1}{Gro_2})$$

Here, $Gro_1$ and $Gro_2$ are the growth rate of one population and its competitor, respectively. Moreover, InifR was introduced to assess the asymmetry of initial abundances of each population, given by

$$\mathrm{InifR} = \log(\frac{Inif_1}{Inif_2})$$

In which, $Inif_1$ and $Inif_2$ are the initial cell numbers of one population and its competitor, respectively. SAR was used to evaluate the asymmetry of manners of colonizing space, which was calculated as described before. Based on these definitions, the values of each parameter greater than 0 denote that the focus population possesses the corresponding competitive edge.

Next, a parameter set containing 89100 combinations of GroR and InifR were designed, in which the values of GroR varied from -0.0513 to 0.0513, and values of InifR were ranged from -2.197 to 2.197. To generate different SAR values, 30 different initial cell distributions were established using the protocol as shown in Supplementary Information S1, which covered a gradient of ScatR values ranging from -0.478 to 0.478. As ScatR was positively correlated with SAR (Fig. S6a), a gradient of SAR values can be observed after performing simulations initialized with these initial distributions. Therefore, 89100 simulations were performed to build the link among these three parameters, in which each simulation was initialized with one of the defined initial distributions, as well as the pre-defined GroR and InifR values. After simulations, SAR values were calculated, ranging from -5.248 to 5.248, and AbunR, as well as final competition outcomes were also calculated and recorded. Two three-dimensional density maps (Fig. 5b and Fig. 5c) were then generated to visualize their relations using the *plot3D* package (v1.3) in R 4.0.2 (https://cran.r-project.org/).

However, higher initial abundance also potentially leads to more seeding positions for the population at the beginning, and hence resulted in an additional advantage from 'Space Accessibility'. To eliminated the effect of initial cell number on the 'Space Accessibility', deriving from the difference in seeding positions at the beginning, a more general parameter perSAR was defined to quantify the asymmetry of 'Space Accessibility' between one population to its competitor, which was calculated by

$$\text{perSAR} = \log\left(\frac{\frac{\sum_{t=0}^{t=t_2} SA_{1,t}}{Inif_1}}{\frac{\sum_{t=0}^{t=t_2} SA_{2,t}}{Inif_2}}\right)$$

Here, $Inif_1$ represents the initial cell number of the population, while $Inif_2$ represents the initial cell number of its competitor. According to this definition, when the population has the same initial cell numbers as its competitor (that is, $Inif_1 = Inif_2$), perSAR equals SAR. When the difference in initial cell numbers is considered, the perSAR indicates the ratio of the 'Space Accessibility' normalized by the initial cell numbers. Analyses about perSAR were performed as the protocol described above.

**S2 Robustness test of the effect of'Space Accessibility'**

In our basic model, we controlled initial conditions to simplify the mathematical analysis. Then, we performed robustness test to investigate whether the effect of 'Space Accessibility' on the outcomes of spatial competition is statistically significant under various initial conditions.

Firstly, to test the effects of the initial number of colonized cells, we performed additional simulations using the same protocol as our basic model except for setting a gradient of the initial total cell number of the two competing populations, ranged from 1% to 50% of the maximum population size (that is, from 4 cells to 200 cells; Table S1), while the initial ratio of the two populations was still set as 1:1. After these simulations, we performed similar correlation analysis as before, and calculated the effect size index (Cohens'D [1]) to assess whether 'Space Accessibility' has a significant impact on the competition outcome under different initial conditions. As shown in Table S1, we found that when the initial total number of the two populations did not exceed to 10% of the maximum population size, 'Space Accessibility' was significantly ($P<0.001$, Cohens'D>0.2) correlated with the competition outcome. In contrast, when the initial cell number was over this threshold, this effect became less significant (Cohens'D<0.2). This shift was attributed to the decreased amount of initial free positions.

Then, we tested whether the effect of 'Space Accessibility' on competition outcome is different when the spatial competition occurred between two faster growing populations, or between two slower growing populations. We changed our basic model settings to consider three different growth rates, 0.01, 0.1 and 1 (Table S1). For each simulation, we assigned the same growth rate to two competing populations and performed correlation analyses after all the simulations. The results showed that the effect of ‘Space Accessibility’ was significant in competition between both faster growing and slower growing populations (Table S1).

Finally, to investigate whether the effect of ‘Space Accessibility’ is still significant when the size of space becomes larger, we simulated two populations competing for space in a larger discrete grid box of a 100×100 array. A new C++ code was written to implement these new simulations (https://github.com/Neina-0830/BacGo-model). Additional simulation results showed that AbunR was still strongly positively correlated with ‘Space Accessibility’ asymmetry SAR (Fig. S6a; $R^2$=0.878, P<0.001). Furthermore, SAR value of the simulations in which the focus population won, was still significantly higher than that for the population lose (Fig. S6b; t-value= -8.028, P< 0.001). These results indicated that larger space size also didn’t change the effect of ‘Space Accessibility’ on the outcomes of spatial competition.

Together, these analyses demonstrated that the effect of ‘Space Accessibility’ on competitive success of a population is robust within a wide range of initial conditions.

### S3 Figures and Tables

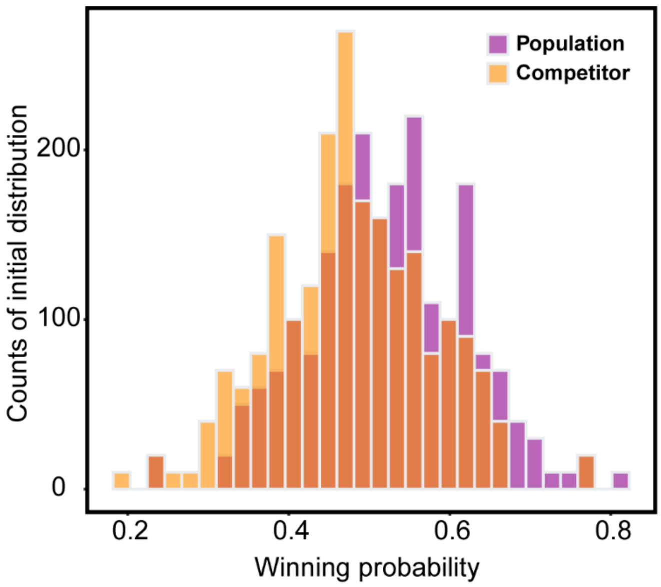


**Fig. S1 Histogram of the frequency distribution of winning probabilities of one population and its competitors.** Results were summarized from 200 random generated initial distributions and each initial distribution was repeated 100 times to calculate the winning probability.

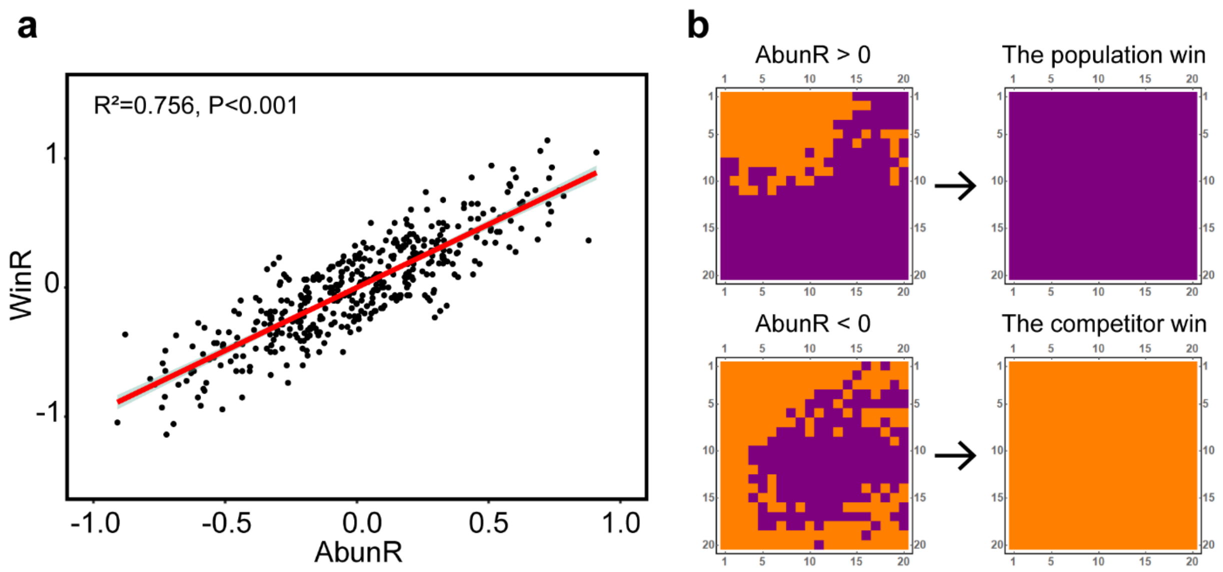


**Fig. S2 Correlation between AbunR and WinR. a.** Correlation between AbunR and WinR. Results represented the sum of 20000 independent simulations, which was same as the Supplementary Figure 1. **b.** Diagram characterizing our hypothesis that the population possessing an AbunR value over zero will possess higher probability to finally win the spatial competition.

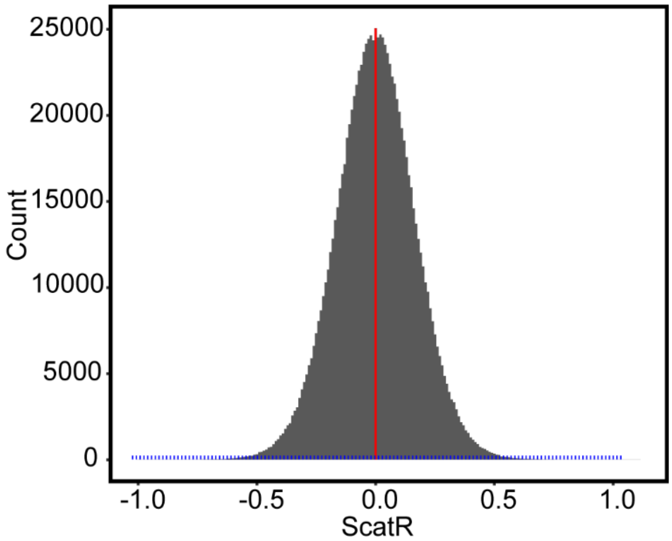


**Fig. S3 Frequency distribution histogram of ScatR with 1000000 randomly generated initial distributions.** The red line is shown at where ScatR equals zero, and the blue lines are evenly distributed over all ranges. 363 initial distributions were selected from the red line to explore the influence of expansion freedom, and 215 initial cell distributions were selected with uniformly gradient ScatR values to explore the influence of initial scattered level, as shown in the blue lines.

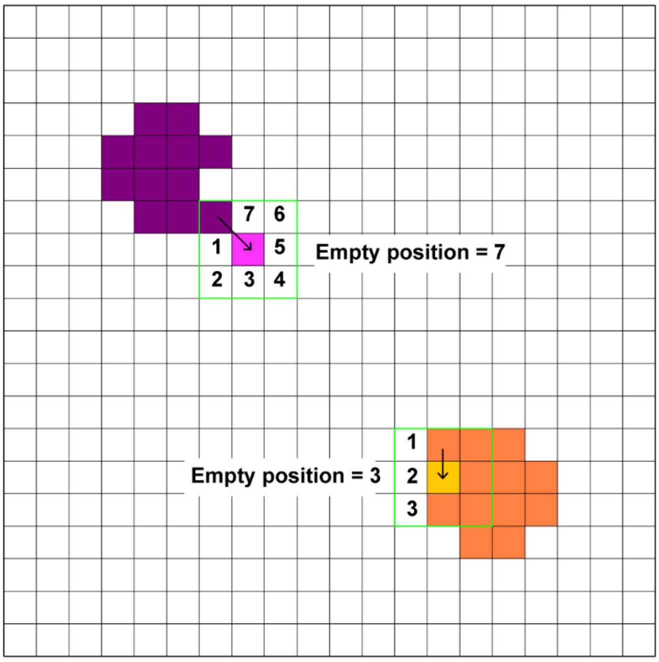


**Fig. S4 Diagram indicating the definition of 'Empty positions' surrounding each newly occupied position.**

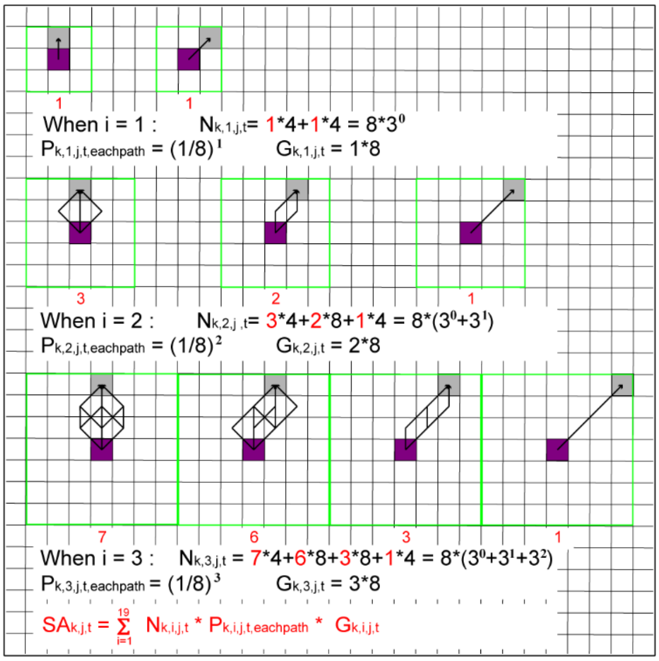


**Fig. S5 Diagrams indicating the calculation methods of $SA_{k,j,t}$.**

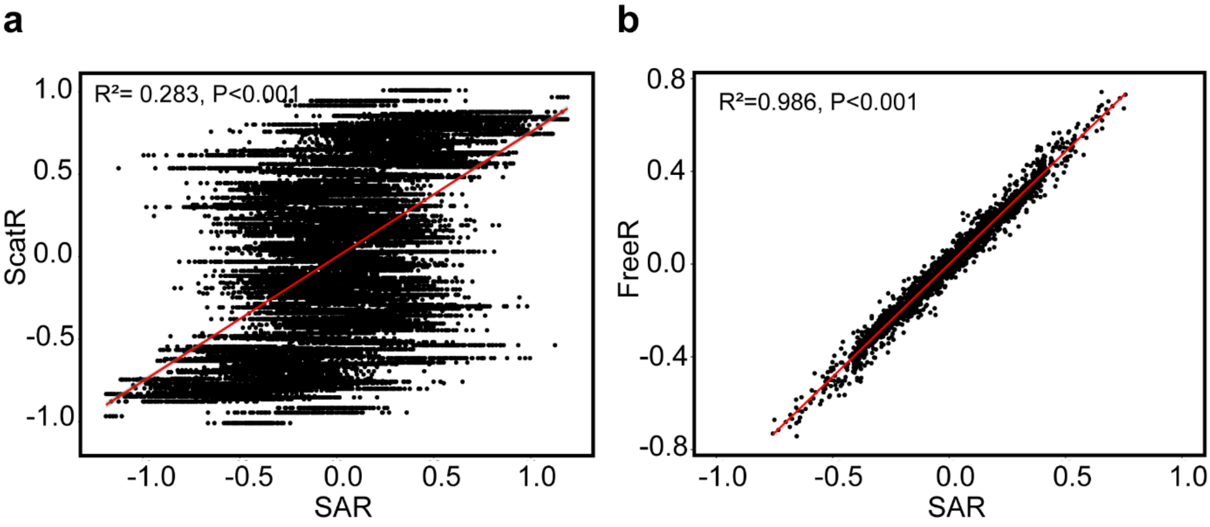


**Fig. S6 Relationship between SAR, ScatR and FreeR. a.** Relationship between SAR and ScatR. **b.** Relationship between SAR and FreeR. Results represented the sum of 20000 independent simulations.

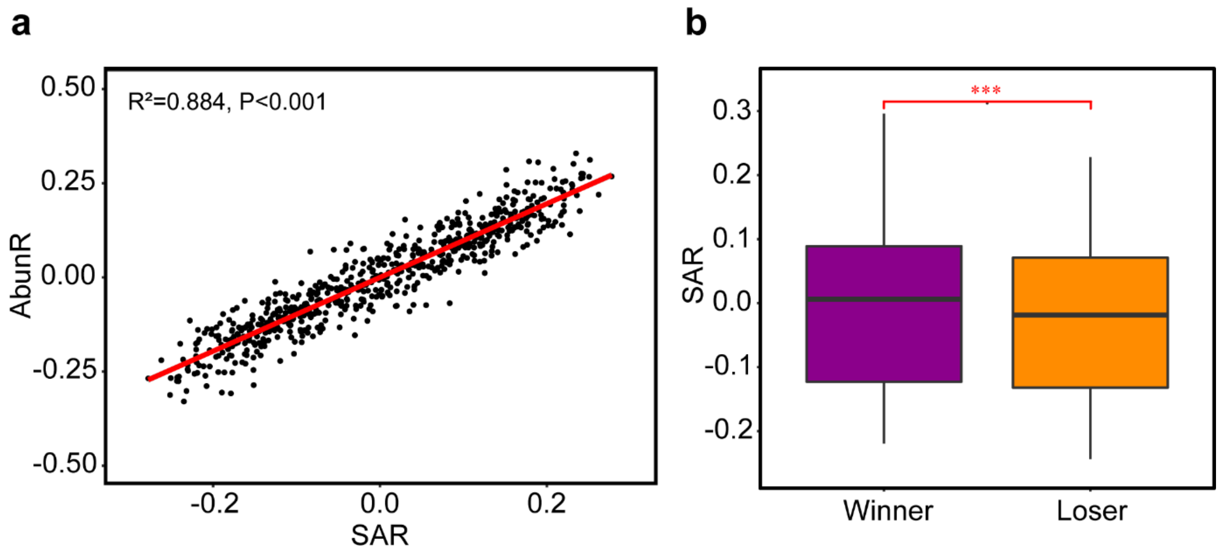


**Fig. S7 Effect of the SAR on the competitive outcomes of a population in a space containing 100×100 array. a.** Correlation between the 'Space Accessibility' asymmetry SAR and the abundance asymmetry AbunR for every population. **b.** Comparation between the 'Space Accessibility' asymmetry of the winning population and that of the failed population. The competition outcomes were generated from same simulations in a. Statistical analysis was performed by two-sample Student's t-test: ***, $P < 0.001$. Results represented the sum of 400 independent simulations.

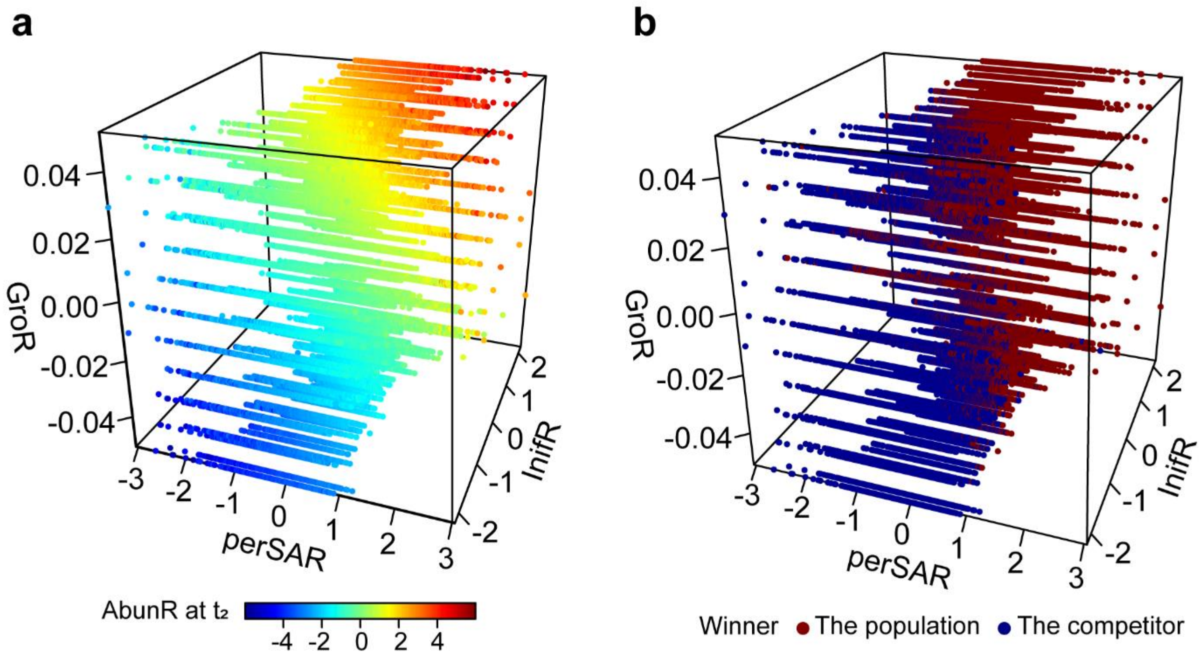


**Fig. S8 Comparison of the relative importance of perSAR, GroR and InifR for the outcomes of microbial competition.** Values of AbunR (**a**), as well as the final competition outcomes (**b**) were also recorded to estimate how these three factors collectively affect the microbial competition.

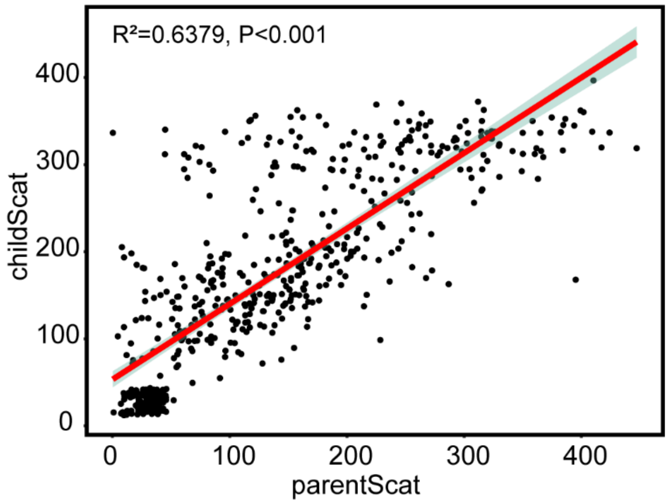


**Fig. S9 Scattered index of child trees positively correlated with that of parent trees.** Data were collected from ForestGEO (https://forestgeo.si.edu/). ParentScat was defined as the average scattered index of these individuals whose DBH were in the top 10% for each tree, and childScat was defined as the average scattered index of other 90% individuals for each tree. All calculations were implemented by Wolfram Mathematica 12.0 (data and codes were available on https://github.com/Neina-0830/BacGo-model).

**Table S1** Robustness test of the effect of 'Space Accessibility' on the competition outcomes.

| **Initial numbers**[a] | **Growth rate** | **Simulation times** | **Determination**[b] | **Effect size**[c] | **P value** |
|---|---|---|---|---|---|
| 100+100 | 0.1 | 1000 | 0.163 | 0.107 | <0.001 |
| 90+90 | 0.1 | 1000 | 0.137 | 0.235 | <0.001 |
| 80+80 | 0.1 | 1000 | 0.253 | 0.141 | <0.001 |
| 70+70 | 0.1 | 1000 | 0.268 | 0.141 | <0.001 |
| 60+60 | 0.1 | 1000 | 0.293 | 0.190 | <0.001 |
| 50+50 | 0.1 | 1000 | 0.314 | 0.062 | <0.001 |
| 40+40 | 0.1 | 1000 | 0.369 | 0.133 | <0.001 |
| 30+30 | 0.1 | 1000 | 0.520 | 0.092 | <0.001 |
| 20+20 | 0.1 | 1000 | 0.630 | 0.238 | <0.001 |
| 10+10 | 0.1 | 1000 | 0.614 | 0.235 | <0.001 |
| 2+2 | 0.1 | 1000 | 0.894 | 0.573 | <0.001 |
| 10+10 | 0.01 | 5000 | 0.716 | 1.022 | <0.001 |
| 10+10 | 0.1 | 5000 | 0.889 | 0.575 | <0.001 |
| 10+10 | 1 | 5000 | 0.850 | 0.431 | <0.001 |

Note:

**a.** Two same numbers in the first column denote the initial cell numbers of two competitive populations.

**b.** The determination coefficient $R^2$ refers to the correlation strength of 'Space Accessibility' asymmetry SAR and abundance asymmetry AbunR at the "full occupied" time ($t_2$).

**c.** Considering that significance level (p-value) is easily affected by sample size, we also calculated the effect size index (Cohens'D [1]) for the purpose of statistical analyses. Only if Cohens'D was greater than 0.2 [2] would SAR be considered to have an impact on AbunR.

**Table S2** Results of multiple linear regression analysis of AbunR with GroR, SAR, and InifR.

| **Parameters** | **Coefficients** | **P-value** | **VIF** | **Adjusted $R^2$** |
|---|---|---|---|---|
| GroR | 55.393 | <0.001 | 1.007 | |
| InifR | 1.027 | <0.001 | 1.008 | 0.993 |
| perSAR | 1.027 | <0.001 | 1.000 | |

**Table S3** Summary of model variables

| Variables | Definition | Units | Values | References |
|---|---|---|---|---|
| $\boldsymbol{B_i}$ | Carbon biomass of the individual in the *i*th grid. | fg | | |
| $\boldsymbol{B_0}$ | Initial C biomass of an individual. | fg | 150 | [3] |
| $\boldsymbol{\mu_i}$ | Growth rate of the individual in the *i*th grid. | fg/fg·min | 0.1 | [3] |
| $\boldsymbol{d_i}$ | Random death rate of the individual in the *i*th grid. | fg/fg·min | $1\times10^{-4}$ | [4] |
| $(\boldsymbol{x_{ki}}, \boldsymbol{y_{ki}})$ | Position coordinate of the *i*th individual of the *k*th population. | | $0 \leq x_{ki}, y_{ki} < 20$ | |

**Table S4** Summary of the defined index

| Defined index | Definition |
|---|---|
| $\boldsymbol{WinR}$ | Asymmetry of winning probability between one population and its competitor. |
| $\boldsymbol{AbunR}$ | Asymmetry of relative abundance at $t_2$ between one population and its competitor. |
| $\boldsymbol{ScatR}$ | Asymmetry of scattered level of the cell distribution at $t_1$ between one population and its competitor. |
| $\boldsymbol{FreeR}$ | Asymmetry of expansion freedom between one population and its competitor. |
| $\boldsymbol{SAR}$ | Asymmetry of 'Space Accessibility' between one population and its competitor. |
| $\boldsymbol{perSAR}$ | The new asymmetry index of 'Space Accessibility' between one population and its competitor. |
| $\boldsymbol{GroR}$ | Asymmetry of growth rate between one population and its competitor. |
| $\boldsymbol{InifR}$ | Asymmetry of relative abundance at $t_1$ between one population and its competitor. |

**Table S5** Summary of symbol

| Symbol | Definition |
|---|---|
| $\boldsymbol{smartBac}$ | Population whose daughter cells always non-randomly select the position to ensure a higher 'Space Accessibility'. |
| $\boldsymbol{normalBac}$ | Population whose daughter cells always randomly select the position. |
| **SA** | 'Space Accessibility' of the population. |
| **Winner** | Population who win the competition. |
| **Loser** | Population whose competitor win the competition. |
| $\boldsymbol{t_1}$ | Time point when the competition is beginning. |
| $\boldsymbol{t_2}$ | Time point when space is full occupied. |
| $\boldsymbol{t_3}$ | Time point when the winner occupies the entire space. |

### SI References


1. J, C. STATISTICAL POWER ANALYSIS FOR THE BEHAVIORAL-SCIENCES Percept. Mot. Skills 67, 1007-1007 (1988).
2. Ma, Z. S., Li, L. & Gotelli, N. J. Diversity-disease relationships and shared species analyses for human microbiome-associated diseases. ISME J 13, 1911-1919, doi:10.1038/s41396-019-0395-y (2019).
3. DK, B. Nutrient Uptake by Microorganisms according to Kinetic Parameters from Theory as Related to Cytoarchitecture. Microbiol Mol Biol Rev 62, 636-645 (1998).
4. Allison, S. D. Cheaters, diffusion and nutrients constrain decomposition by microbial enzymes in spatially structured environments. Ecology Letters 8, 626-635, doi:10.1111/j.1461-0248.2005.00756.x (2005).
